## Supplementary Information for "High-Resolution Proteomics Unveils Salivary Gland Disruption and Saliva-Hemolymph Protein Exchange in *Plasmodium*-Infected Mosquitoes"

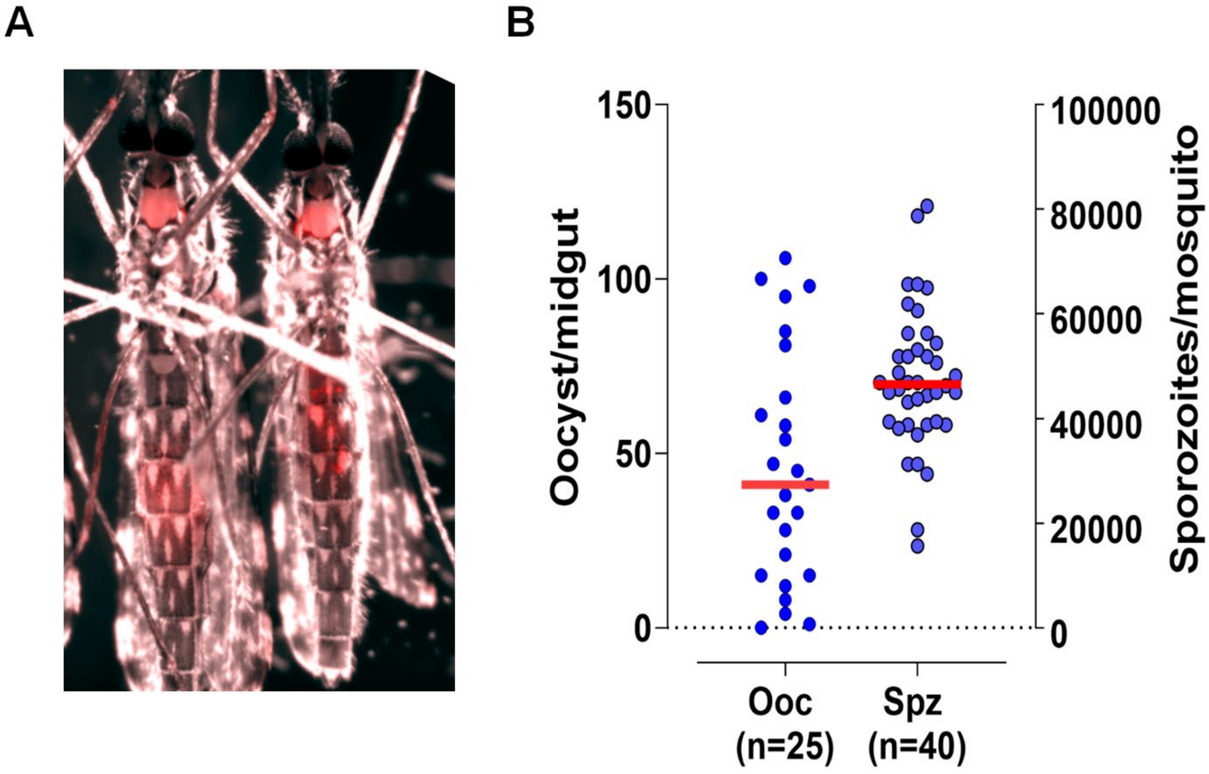


**Supplementary Figure 1: Sorting *P.berghei* mCherry-infected *An. gambiae* mosquitoes.** (A) Infected mosquitoes were sorted by fluorescence (GFP or mCherry) with a Leica M205 fluorescence stereo microscope FCA coupled with a DFC 7000 G5 camera. (B) Distribution of oocysts and sporozoites in a representative proteome experiment.

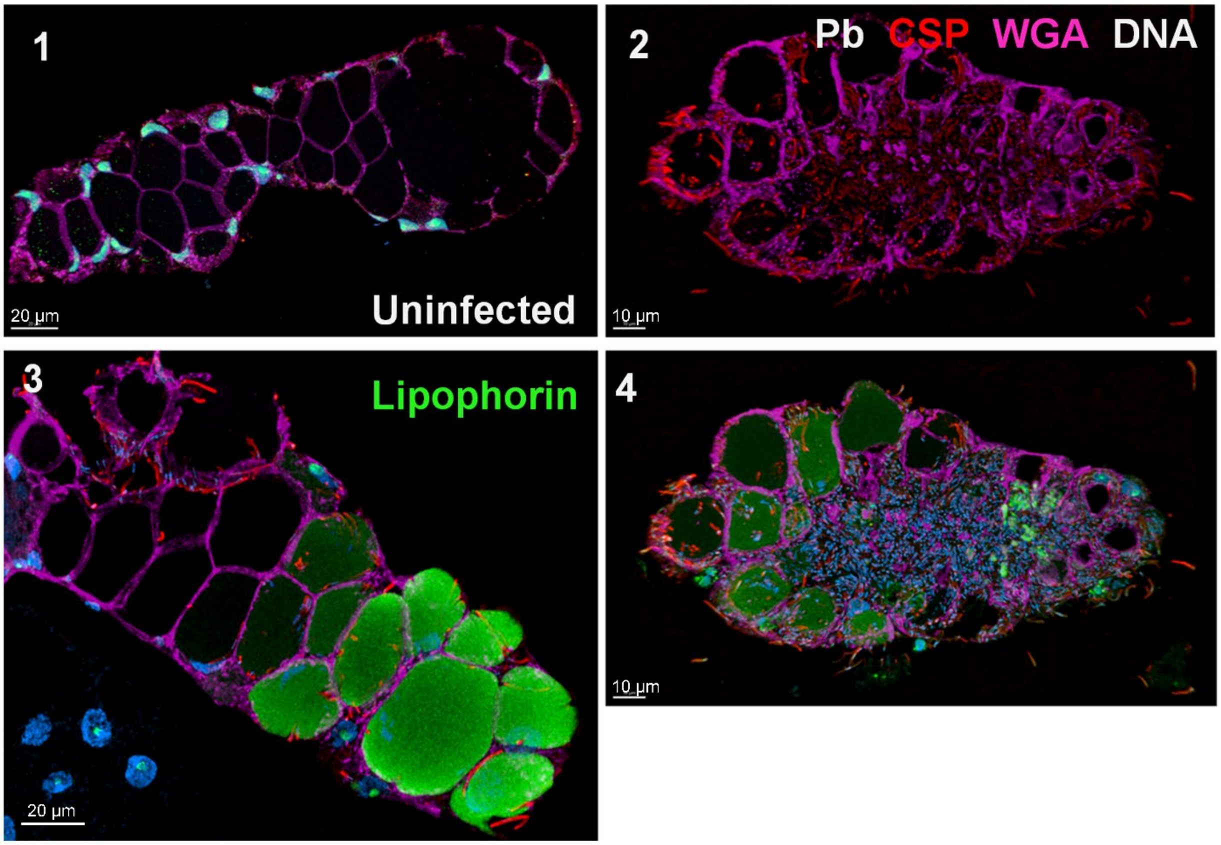


**Supplementary Figure 2: Immunofluorescence assay of lipophorin accumulation in infected salivary glands.** Staining: Lipophorin green, CSP red, DNA blue, WGA magenta. 1 uninfected, 2-4 *P. berghei* infected salivary glands. Infected salivary glands show increased accumulation of lipophorin.

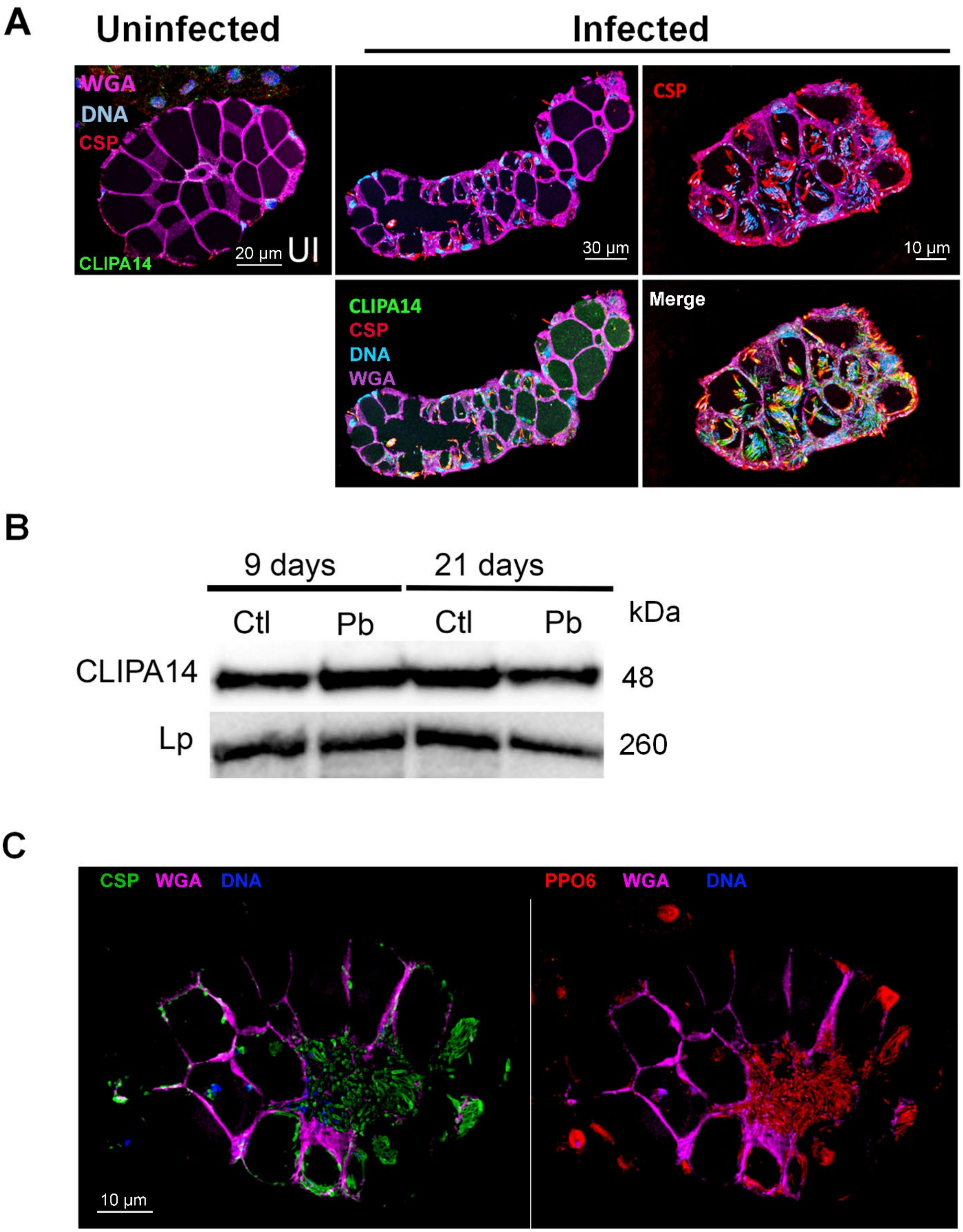


**Supplementary Figure 3: Histological immunofluorescence assay of uninfected and *P. berghei* infected salivary glands.** (A) CLIPA14 green, CS red,

WGA magenta, DNA blue. (B) Western Blot Analysis of CLIPA14 Expression: Western blot showing CLIPA14 expression in uninfected (UI) and *P. berghei*-infected (Pb) hemolymph at 9 and 21 days post-infection. At 9 days, no invasion of salivary glands is detected, whereas at 21 days, the salivary glands are heavily infected. After apyrase staining (Fig. 5C), the membrane was stripped and reprobed for CLIPA14. Lipophorin was used as a loading control. (C) PPO6 red, CS green, WGA magenta, DNA blue.

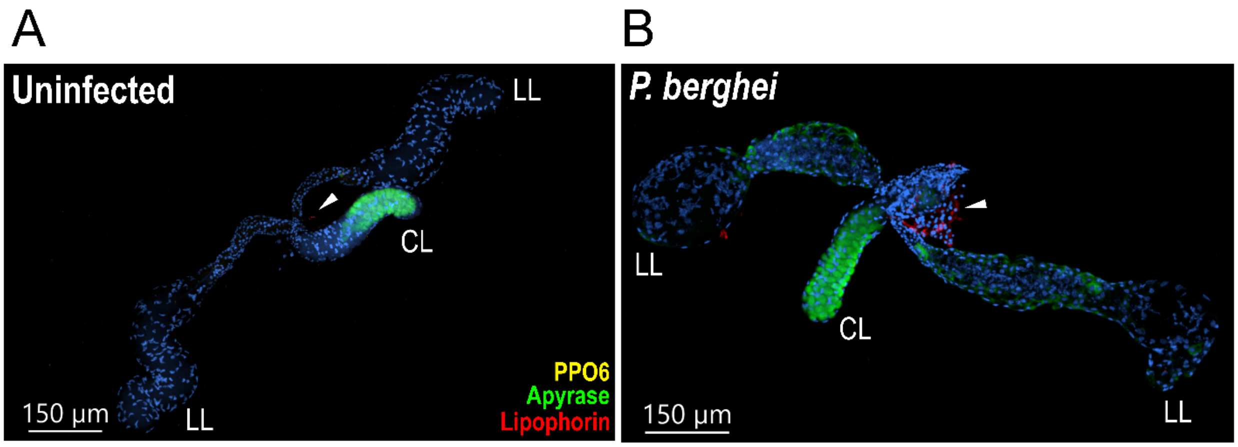


**Supplementary Figure 4. Additional examples of the transcriptional activity in uSGs (U) and iSGs (Pb).** RNA in situ hybridization in (A) uSGs and (B) iSGs. Nuclei in blue, PPO6 in yellow, Apyrase in green, and Lipophorin in red. (LL) lateral lobe and (CL) central lobe. The arrow in (A) and (B) indicate fat body cells associated with SGs.

**A** **Most abundant hemolymph proteins from *P. berghei*-infected mosquitoes 19 days post-infection. Bolded proteins enriched in iSGs.**

| Protein IDs | Description |
|---|---|
| AGAP004977-PA | **Prophenoloxidase 6** |
| AGAP001826-PA | **Lipophorin** |
| AGAP006258-PA | Prophenoloxidase 2 |
| AGAP011369-PA | **Gelsolin** |
| AGAP000376-PA | Transferrin |
| AGAP011788-PA | **CLIPA14: CLIP-domain serine protease** |
| AGAP003250-PA | **CLIPB4: CLIP-domain serine protease** |
| AGAP010968-PA | CLIPA9: CLIP-domain serine protease |
| AGAP003095-PA | **Yellow protein** |
| AGAP011789-PA | **CLIPA6: CLIP-domain serine protease** |
| AGAP008060-PA | **IDGF2: imaginal disc growth factor 2** |
| AGAP008364-PG | **TEP15: thioester-containing protein 15** |
| AGAP011792-PA | **CLIPA7: CLIP-domain serine protease** |
| AGAP003139-PA | Serine protease inhibitor (serpin) 9 |
| AGAP008061-PA | **IDGF4: imaginal disc growth factor 4** |
| AGAP012616-PA | Prophenoloxidase 5 |
| AGAP004976.P46 | Prophenoloxidase 8 |
| AGAP004978-PA | Prophenoloxidase 9 |
| AGAP010658-PA | Hexamerin |
| AGAP001377-PA | **Serine protease inhibitor (serpin) 11** |
| AGAP002925-PC | **Poly(U)-specific endoribonuclease** |
| AGAP004455-PA | GNBPB1: 3-glucan binding protein |
| AGAP002465-PA | Ferritin heavy chain |
| AGAP001662-PA | Disintegrin metalloproteinases with thrombospondin repeats |
| AGAP029749-PB | Trypsin-like serine protease |
| AGAP002032-PA | Lipoprotein |
| AGAP007455-PA | LRIM10: Leucine-rich immune protein (Short) |
| AGAP008364-PD | **TEP15: Thioester-containing protein 15** |
| AGAP009670-PC | SRPN4: Serine protease inhibitor (serpin) 4 |
| AGAP006278-PA | D7 protein |

**B**

### SGs proteome: Proteins up-regulated in iSGs. Abundant hemolymph proteins are bolded

| Protein IDs | Description | Fold change |
|---|---|---|
| AGAP001569-PB | Myosin Alkali Light Chain 1 | 17.4854256 |
| AGAP010935-PA | Porphobilinogen Synthase | 14.1053903 |
| **AGAP003250-PA** | **CLIPB4** | 12.7574957 |
| AGAP001023-PF | Myofilin Variant B | 10.6730819 |
| AGAP002350-PB | Troponin T, Fast Skeletal Muscle | 10.4584351 |
| **AGAP002925-PC** | **Poly(U)-Specific Endoribonuclease** | 9.76303695 |
| **AGAP011788-PA** | **CLIPA14** | 6.87116899 |
| AGAP011369-PA | Gelsolin | 6.5154338 |
| **AGAP000376-PA** | **Transferrin 1** | 4.82313294 |
| AGAP012401-PA | AGM1 | 3.83590822 |
| AGAP012115-PA | $Ca^{2+}$-Transporting ATPase | 3.50907114 |
| AGAP001826-PA | **Lipophorin** | 3.48664891 |
| **AGAP008060-PA** | **Imaginal Disc Growth Factor 2** | 3.09660848 |
| AGAP005712-PC | Phenylalanine-4-Hydroxylase | 2.87706371 |
| **AGAP004977-PA** | **PPO6** | 2.44134122 |
| AGAP006103-PA | AGAP006103-PA | 2.27812436 |
| **AGAP008061-PA** | **IDGF4: Imaginal Disc Growth Factor 4** | 2.22152879 |
| **AGAP008364-PA** | **TEP15** | 2.16172823 |
| AGAP011842-PA | Signal Peptidase Complex Subunit 2 | 2.09807517 |
| AGAP004960-PA | Prosalpha3: 26S Proteasome Alpha 3 Subunit | 1.93335732 |
| AGAP001271-PA | Pre-mRNA Cleavage Complex 2 Protein (Pcf11) | 1.91235929 |
| AGAP002335-PB | Nucleolysin TIA-1/TIAR | 1.89223474 |
| AGAP001313-PA | Muscular Protein 20 | 1.85127917 |
| AGAP008774-PA | Cytochrome C Oxidase Assembly Protein | 1.65646615 |

### SGs proteome: Proteins unique to iSGs. Abundant hemolymph proteins are bolded

| Protein IDs | Description |
|---|---|
| AGAP005625-PA | SCRASP1: Class A Scavenger Receptor |
| AGAP003865-PA | gamma-tubulin complex component 3 |
| **AGAP011789-PA** | **CLIPA6** |
| **AGAP011792-PA** | **CLIPA7** |
| AGAP007249-PB | Flightin: protein flightin |
| AGAP001905-PB | zinc finger RNA-binding protein |

**C**

**Hemolymph Proteome of Infected Mosquitoes 19 Days Post-Infection: Salivary Proteins Detected Exclusively in Infected Samples. Bolded proteins indicate those with spectral counts present in both replicates**

| | | | *P. berghei* | | Uninfected | |
|---|---|---|---|---|---|---|
| Protein ID | Description | Unique peptides | MS/MS count | MS/MS count | MS/MS count | MS/MS count |
| **AGAP011026-PA** | Apyrase | 14 | 2 | 18 | 0 | 0 |
| **AGAP006421-PA** | Antigen 5 Related Protein 1 | 10 | 3 | 23 | 0 | 0 |
| AGAP008282-PA | D7r2 | 3 | 1 | 4 | 0 | 0 |
| AGAP009917-PA | SGS4 | 68 | 0 | 108 | 0 | 0 |
| AGAP009918-PA | SGS5 | 57 | 0 | 69 | 0 | 0 |
| AGAP008279-PA | D7L2 | 5 | 0 | 7 | 0 | 0 |
| AGAP008278-PA | D7L1 | 4 | 0 | 4 | 0 | 0 |
| AGAP008281-PA | D7r4 | 4 | 0 | 4 | 0 | 0 |
| AGAP009974-PA | Anopheline Antiplatelet Protein | 2 | 0 | 2 | 0 | 0 |
| AGAP008284-PA | D7r1 | 2 | 0 | 2 | 0 | 0 |
| AGAP008283-PA | D7r3 | 2 | 0 | 2 | 0 | 0 |

**Supplementary Figure 5. Most abundant proteins from proteome analysis of hemolymph, salivary glands, and saliva.** (A) List of the most abundant secreted proteins detected by proteomic analysis of hemolymph from *P. berghei*-infected mosquitoes 19 days post-infection. The proteins are ranked from the most abundant to the least. Bolded names are proteins found enriched in infected salivary glands. (B) Proteomic analysis of SGs from uninfected and *P. berghei*-infected mosquitoes. The table lists proteins upregulated or enriched in infected samples and their respective fold changes. Bolded are putative hemolymph proteins enriched in infected salivary glands. The upper table shows proteins found in both uninfected and infected samples, while the lower table lists proteins detected exclusively in infected samples. Proteins abundant in the mosquito hemolymph are bolded. (C) Saliva proteins detected exclusively in the hemolymph of *P. berghei*-infected mosquitoes.

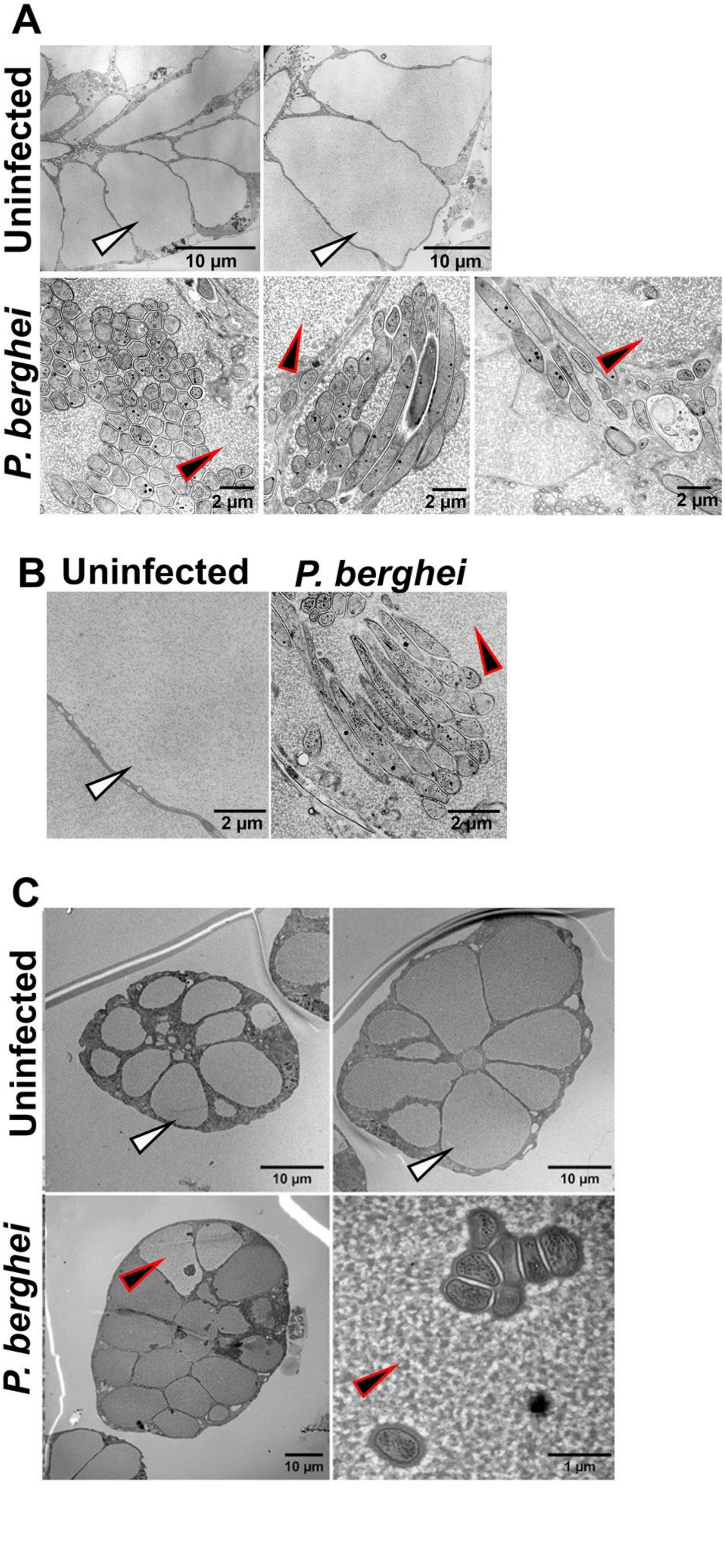
A
Uninfected
10 µm
10 µm
P. berghei
2 µm
2 µm
2 µm
B
Uninfected
P. berghei
2 µm
2 µm
C
Uninfected
10 µm
10 µm
P. berghei
10 µm
1 µm

**Supplementary Figure 6: *Plasmodium berghei* infection of salivary glands changes the texture of saliva within the secretory cavities.** (A) Uninfected and *P. berghei* ANKA 2.34-infected SGs from *Anopheles stephensi*. (B) A closer view at the same scale of uninfected and *P. berghei*-infected *An. stephensi* SGs. Images in panels C and D were generated in Dr. Isabelle Coppens' lab at Johns Hopkins University. (C) *An. stephensi* SGs infected with *P. berghei* ANKA 2.34. The arrow points to an infected cavity with distinct granularity in the saliva matrix. The electrolucent and granular aspects of the saliva are more evident in the bottom right. These images were generated in Dr. Friedrich Frischknecht's lab at Heidelberg University Medical School.

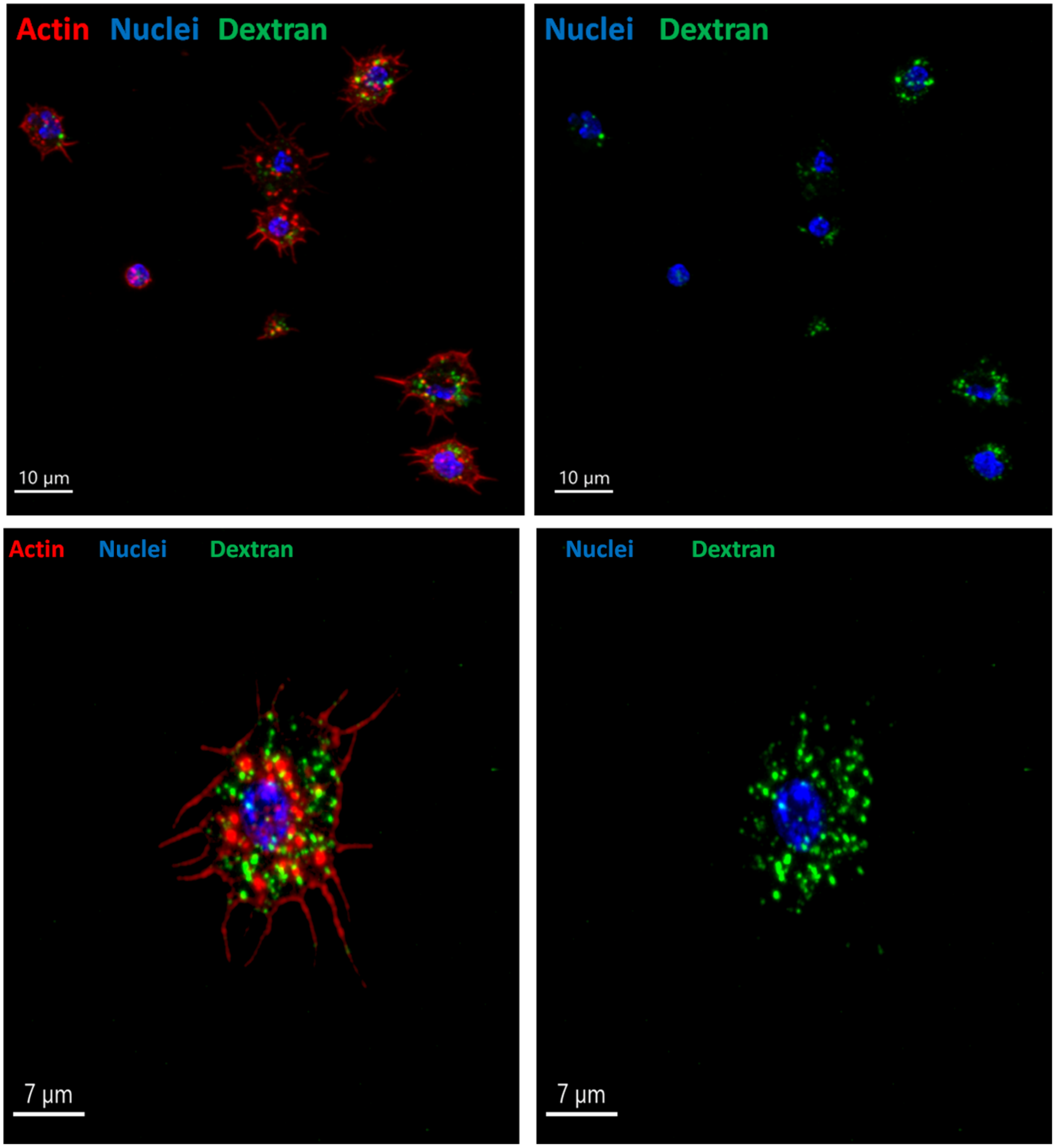


**Supplementary Figure 7. Hemocytes capture injected dextran in mosquitoes.** Fluorescent dextran was injected into mosquitoes, and hemocytes were subsequently collected by perfusing the mosquito hemolymph. Hemocytes are shown with actin (phalloidin) in red, dextran in green, and nuclei (Hoechst) in blue.

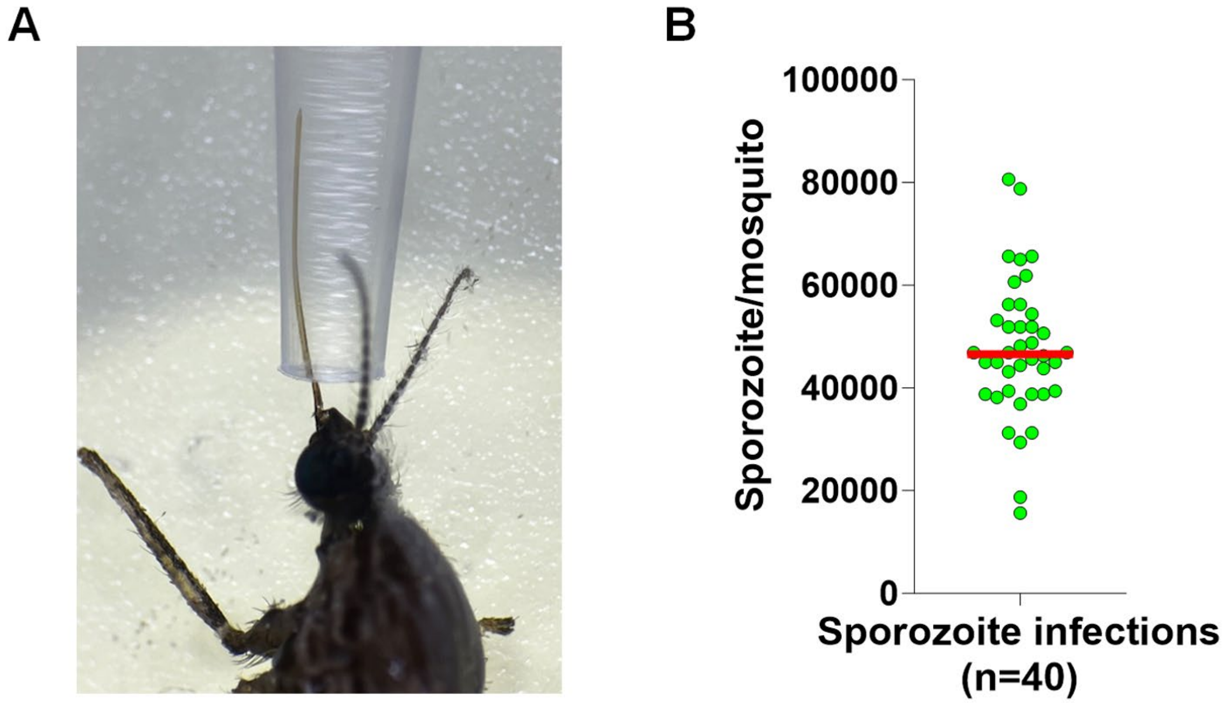


**Supplementary Figure 8.** (A) Collection of mosquito saliva. (B) Representative *P. berghei* sporozoite counting in infected salivary glands.

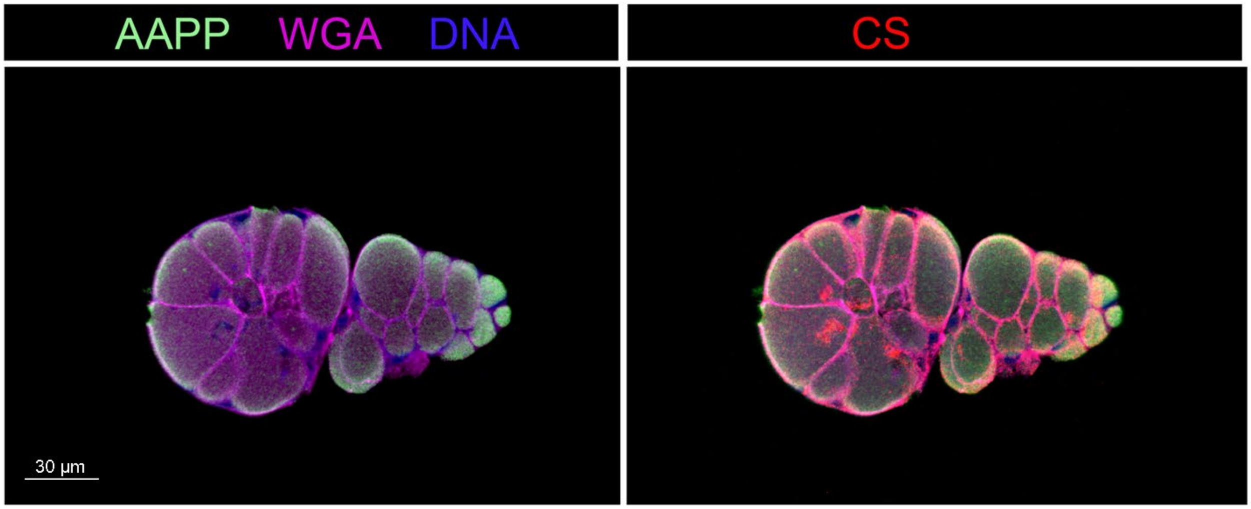


**Supplementary Figure 9. Histological immunofluorescence of *P.berghei*-infected salivary glands.** These images display the distribution of AAPP within the infected salivary gland. Staining includes AAPP (green), circunsporozoite (red), and WGA (magenta).
